# The Influence of Leaf Anatomical Traits on Photosynthetic in Coffee Leaves

**DOI:** 10.1101/2022.02.18.481015

**Authors:** Ziwei Xiao, Guanrun Ma, Xuehui Bai, Jinhong Li, Mingzhu Zhao, Linlin Su, Hua Zhou

## Abstract

Leaf photosynthesis is largely determined by anatomical features. This study aimed to reveal the quantitative effects of the anatomical structure of *Coffea arabica* leaves on photosynthesis. Pearson’s correlation and path analysis were used to explore the correlation between leaf structure and photosynthesis. To calculate the comprehensive evaluation value of the correlation between leaf anatomical traits and photosynthetic parameters, the Criteria Importance Through Intercriteria Correlation (CRITIC) method was used to obtain the objective weight of each photosynthetic parameter. Most of the anatomical traits of the leaf significantly (*P*<0.05) to highly significantly (*P*<0.01) correlated with photosynthetic parameters, suggesting that anatomical traits greatly affected the photosynthetic capacity in *Coffea arabica* leaf. Likewise, path coefficient analysis revealed that the photosynthetic capacity of Arabic coffee leaves is strongly correlated, both directly and indirectly, with the stomatal characteristics of the leaves. The results of the comprehensive evaluation also indicated that leaf thickness (LT) and stomatal density (*d*) were the anatomical characteristics most closely related to photosynthesis. In these results, understanding the effects of the anatomical structure of coffee leaves on photosynthesis, may provide useful information for coffee breeding programs and the management of coffee plantations to increase photosynthesis.

## Introduction

Coffee (*Coffea arabica*) considered an important tropical crop, is one of the most traded agricultural products in the world. Under suboptimal growing conditions, shade trees, which could reduce the stress on coffee by ameliorating adverse climatic conditions, have been recommended (Beer et al. 2001; Franck and Vaast 2009a). Yunnan province, located in Southwest China, is a predominantly coffee-growing region (ICO 2015). Although the mountainous landscape and mild climate in this region are well-suited to coffee production, many areas are suboptimal for coffee cultivation (ICO 2015; Ma et al. 2019). Thus, agroforestry systems have been recommended for these areas, especially at altitudes below 1000 m (Ma et al. 2019).

Because leaves are specialized organs that enable plants to intercept light and uptake CO_2_ through the stomata for photosynthesis (Lambers and Oliveira 2019), light and CO_2_ availability in the mesophyll of plant leaves largely determines the net carbon assimilation rate (A) of leaves (Franck and Vaast 2009b). To analyze the response of photosynthesis to irradiance, the light response curves (LRCs) were used to describe the relationship between photosynthesis and light intensity (Herrmann et al. 2020). It is an important method to elucidate the response mechanism of photosynthesis and evaluate photosynthetic efficiency by fitting of light-response model (Lambers and Oliveira 2019). Often, LRCs are mathematically described using empirical models. The photosynthetic parameters, such as the information of the maximum photosynthetic capacity (*P*_nmax_), intrinsic and apparent quantum yield (IQY and AQY), light compensation and saturation point (LCP and LSP), and dark respiration rate (*R*_D_), can be obtained from the LRCs. In addition, mechanistic information about the key biochemical parameters of carbon assimilation can be estimated by fitting *A/C*i curves with the mechanistic Farquhar-von Caemmerer-Berry (FvCB) biochemical model described by Farquhar et al. (1980) (Farquhar et al. 1980; Sun et al. 2014; Wang et al. 2017). This model has been widely used to simulate CO_2_ assimilation and applied for decades (Farquhar et al. 1980; GU et al. 2010). It elegantly explains the physiological properties of photosynthesis of intact leaves (Lambers and Oliveira 2019), and the parameters reflecting the biochemical properties of leaves were easier to estimate from this model with gas exchange measurements (Wang et al. 2017).

The structural organization of plant leaves could facilitate or sometimes constrain the photochemical reactions of the photosynthetic process (Boom et al. 2012; Oguchi et al. 2018). Therefore, the leave structural changes may affect photosynthetic efficiency (Akhkha et al. 2001; Herrmann et al. 2020). However, although a variety of ecological studies have revealed a correlation between leaf structural parameters and photosynthetic performance (Bolhar-Nordenkampf and Draxler 1993; James and Bell 2001; Vogelman et al. 1996), there were contradictory and not universal differences between foliage structural and functional variables in the previous work (Niinemets 1999). To understand the effect of leaf structure on each of the photosynthetic performance parameters, it is necessary to analyze the relationship between leaf structure and leaf photosynthesis and to comprehensively evaluate the importance of each component in leaf photosynthesis. Furthermore, evaluation of the relationship between leaf structure and leaf photosynthesis could provide ecophysiological evidence for understanding the structural properties of leaves and how they affect photosynthetic performance. Even though a number of studies analyze the effects of leaf structure on photosynthetic performance (Bolhar-Nordenkampf and Draxler 1993; Oguchi et al. 2018), there are few works synthetically analyzing the correlation between leaf structural parameters and photosynthetic performance (Boom et al. 2012). In particular, little research studies related to coffee were reported. Therefore, this study aimed to explore the relationship between leaf structural parameters and photosynthetic performance, and clarify the importance of the correlation between leaf structural parameters and photosynthetic performance by using a comprehensive evaluation method. Obtaining these results may provide useful information for future coffee breeding and plantation management.

## 2 Materials and methods

### 2.1 Study site characterization and plant material

The experiment was conducted in the shade coffee plantations of Dehong Tropical Agriculture Research Institute of Yunnan (DTARI), Ruili, southwest China. The plantations are located at 890 meters of altitude, at 24.025 south latitude and 97.855west longitudes, with an average temperature of 22.2 °C and precipitation of 1260 mm in the studies conducted years (2019-2021). The soil type in the plantation is acidic lateritic red soil.

The coffee trees of the cultivar are ‘Catimor’, an interspecies hybrid derived from the cross between Caturra (*Coffea arabica* L.) and Timor (*C. arabica* × *C. canephora* Pierre ex Froehner). The coffee trees were approximately five years old and spaced 1×2 m. Shade trees (*Dimocarpus longan* Lour.) planted at 12×4 m, East to West rows-oriented.

Fourteen plants with consistent growth were selected in the coffee plantation, and three leaves were selected on each plant in different directions to measure the leaf gas exchange parameters. All the measurement leaves used the third or fourth completely expanded leaf pair from the apex of the plagiotropic (lateral) branches, located in the upper third of the plant.

### 2.2 Gas exchange measurements

The leaf gas exchange parameters (net CO_2_ assimilation rate (A), stomatal conductance to water vapor (*g*_s_), transpiration rate (E), internal CO_2_ concentration (*C*_i_), vapor pressure deficit (VPD), and leaf water use efficiency (WUE=A·E^-1^) and environmental indicators (leaf temperature (TL), relative humidity (RH) and ambient temperature (TA)) were determined simultaneously by using a portable photosynthesis system CIRAS 3 (PP Systems, USA). Leaf water use efficiency (WUE), representing the units assimilated of CO_2_ per unit of water lost through transpiration, was determined via user manual version 1.06. All measurements were taken on sunny days from 09:00 to 11:30 and 15:00 to 18:00. The reference CO_2_ concentration was set at 390 ppm using a buffer bottle, the temperature at 27 °C, flow rate at 300 μmol.s^−1^, relative humidity at 50 ± 10%, and irradiance at 800μmol m ^−2^.s^−1^. The leaf photosynthetic light response (A/PPFD) and internal CO_2_ response (A/C_i_) curves were constructed by using leaf gas exchange measurement. To collect data for building A/PPFD curves, net photosynthesis was measured at 0, 50, 100, 150, 200, 250, 300,350, 400, 500, 600, 700, 800, 1000, and 1200 μmol m^−2^.s^−1^ (PAR) with a cuvette environment that maintained leaf temperature at 27 °C, reference CO _2_ concentration at 400 ppm. The gas exchange rate usually stabilizes within approximately 3-4 min after the leaf tissue is clamped in the leaf chamber. Data for the construction of the A/Ci curves were collected immediately following the A/PPFD measurements from the same sample leaf. *P*_*n*_ was measured at CO_2_ concentrations of 400, 300, 200, 100, 150, 250, 350, 450, 550, 650, 750, 850, 950, 1050, 1250, 1450 and 1650 μmol.m^-2^.s^-1^ at photosynthesis saturating irradiance 800μmol m^−2^.s^−1^,which light intensity is sufficiently high to saturate the photosynthetic machinery without causing photoinhibition. A total of 41 A/PPFD and 42 A/C_i_ curves were generated.

### 2.3 Measurement of leaf anatomical traits

Following the gas exchange measurement, five samples from leaves of each plant were harvested, sealed in polyethylene bags, and immediately brought back to the laboratory. Two segments (3 mm× 5 mm) were cut from each leaf halfway between the leaf apex and base. The material was then dehydrated on sucrose solutions, followed by the inclusion of Gum Arabic aqueous solution. Leaf transverse sections (7–10 μm) thickness cut using a rotary microtome (Leica CM3050S, Germany) and stained with toluidine blue. Surface sections of the leaves were taken manually and stained with safranine aqueous solution (0.1%). For each leaf, five transversal sections and surface sections were selected systematically to measure the leaf thickness (LT), upper cuticle thickness (UCT), upper epidermal thickness (UET), palisade tissue thickness (PT), spongy tissue thickness (ST), lower epidermal thickness (LET), lower cuticle thickness (LCT), stomatal density (*d*), and guard cell length (*l*) and width (*w*), using a microscope binocular with 10 × and 40 × objectives (Leica DM5000 B). Subsequently, five photographs of microscope fields per section were taken with a photomicroscope equipped with a Leica DFC500 camera (Leica Microsystems, Rijswijk, The Netherlands). The images were analyzed with the Leica LAS X software. The total stomach pore area (TSP) was calculated as *d* ×1/4π × *l* × *w* (7), and by assuming that the stomatal pore was an ellipse(Batos et al. 2010; James and Bell 2001; Patricia et al. 2014). Here *d* is the stomatal frequency calculated by dividing the stomatal count by the area of the field of view, *l* is the guard cell length, and *w* is the guard cell width.

### 2.4 Calculations and statistical analysis

A/PPFD curves were fitted by applying a modified rectangular hyperbola model using Photosynthetic Calculation Software 4.1.1(Ye 2007; Ye 2010; Ye and Yu 2008). The modified rectangular hyperbola model is described by Equation (1).

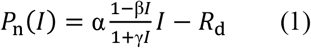

where *P*_*n*_ is the net photosynthetic rate (μmol·CO_2_.m^-2^·s ^-1^), *I* is the photosynthetic photon flux density (μmol·m^-2^·s ^-1^), *I*_c_ is compensation irradiance, β and γ are the coefficients which are independent of *I*, and *α* is the absolute value of slope between *I* = 0 and *I* = *I*_c_.

For *I* = 0, the quantum yield at this point is defined as intrinsic quantum yield (IQY), which is given by Equation (2).

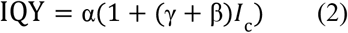

The apparent quantum yield (AQY) ignored the Kok effect is obtained by Equation (3).

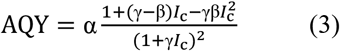

For *I* = 0, the rate of dark respiration (*R*_d_) is Equation (4).

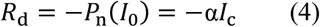

The saturation irradiance *I*_sat_ is obtained by Equation (5).

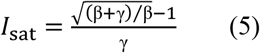

The maximum photosynthetic rate *P*_nmax_ is given by Equation (6).

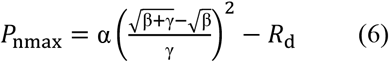

A/Ci curves were fit by uploading the datasets to the website (http://Leafweb.ornl.gov)to estimate the parameters (*R*_d_, *V*_cmax_, *J*_max_, TPU, *Γ*^*^, *g*_i_). In this study, the A/C_i_ curves were taken at the saturation light level. Thus, the rate of electron transport (*J*) was to assume the maximum rate of electron transport (*J*_max_).

The relationship between photosynthetic capacity and leaf anatomical traits was analyzed by calculating the Pearson correlation coefficient. The correlation coefficients were partitioned into direct and indirect effects using path analysis. The path diagram in Figure 1 was used to examine the causal path of nine selected leaf anatomical traits to each photosynthetic parameter. In Figure 1, the direct effects of leaf anatomical traits on photosynthetic parameters are represented by single-headed arrows, while coefficients of intercorrelations between leaf anatomical traits are represented by double-headed arrows. Path coefficient analysis was calculated as the method described by Dewey and Lu (1959) (Dewey and Lu 1959).

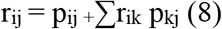

where r_ij_ represents the simple correlation coefficient between the independent character (i) and dependent character (j) as measured by the correlation coefficient, P_ij_ is the component of direct effects of the independent character (i) and dependent character (j) as measured by the path coefficient and, ∑r_ik_ p_kj_ is the summation of components of the indirect effect of a given independent character (i) on the given dependent character (j) via all other independent characters (k). An uncorrelated residue (U) was estimated by using the Equation (9).

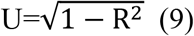

where R^2^ is the coefficient of determination, R^2^ =∑r_ij_ p_ij_.

**Fig 1.**
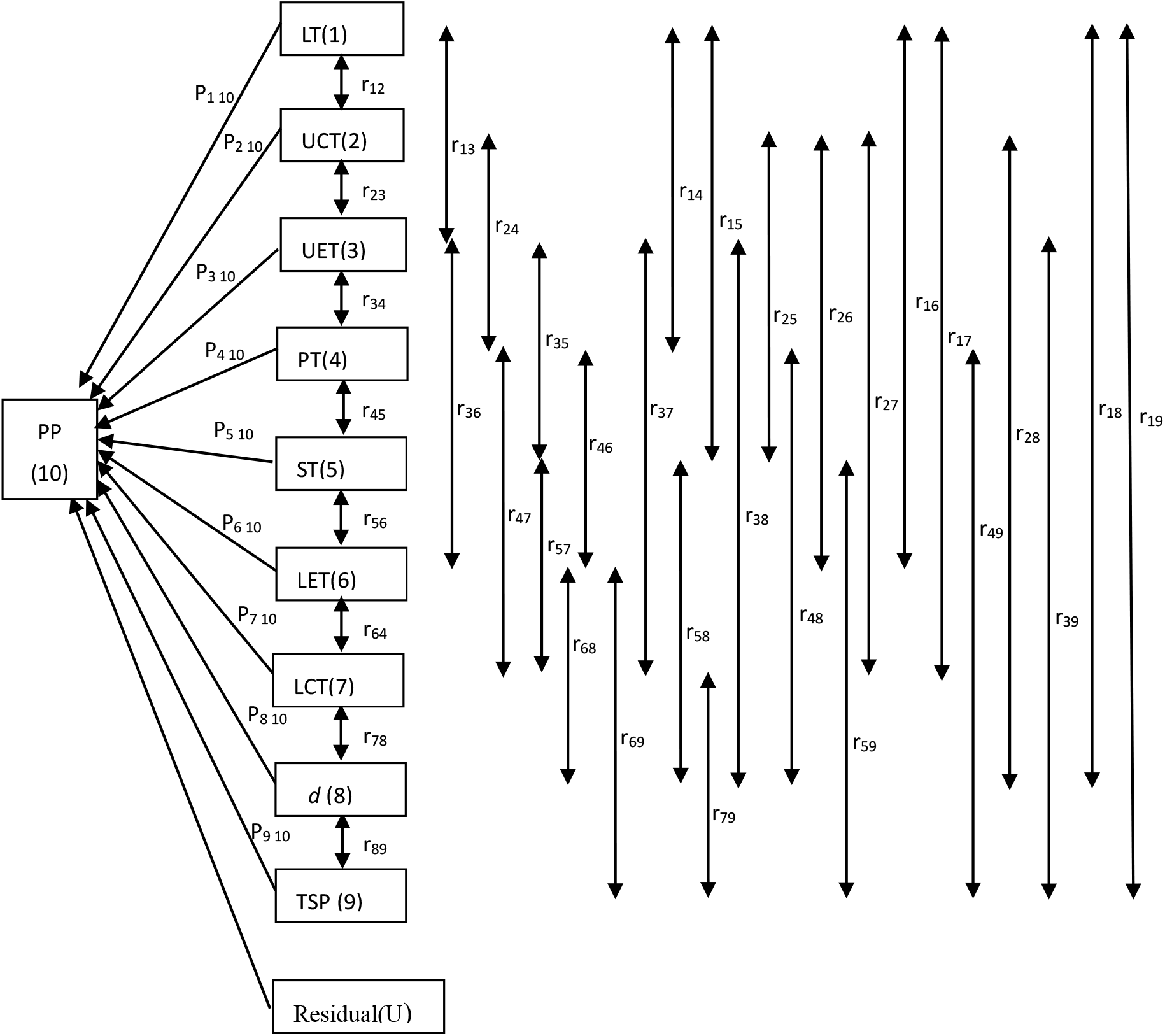
Path diagram for the relationship between leaf anatomical traits and photosynthetic parameters. The direct effects (*P*_*ij*_) of leaf anatomical traits on the photosynthetic parameter (PP) are represented by single-headed arrows, while the indirect effects (*r*_*ij*_*P*_*ij*)_ of leaf anatomical traits are shown by double-headed arrows. Subscript designations for leaf anatomical traits and photosynthetic parameters are identified numerically as follows: (1) LT = leaf thickness; (2) UCT = upper cuticle thickness; (3) UET = upper epidermal thickness; (4) PT = palisade tissue thickness; (5) ST = spongy tissue thickness; (6) LET = lower epidermal thickness; (7) LCT = lower cuticle thickness; (9) *d* = stomatal density; (10) TSP = total stomach pore area; and (10) PP = photosynthetic parameters. The photosynthetic parameters (PP) are IQY, AQY, *α, P*_max_, *I*_sat_,*I*_c_, *R*_D_, *R*_d_, *V*_cmax_, *J*_max_, TPU, *Γ* ^*^, *g*_i_, C_i_, *g*_s_, VPD, A, E, WUE.

The multiple criteria decision analysis (MCDA) was used to evaluate comprehensive evaluation of the relationship between leaf anatomical traits and photosynthetic parameters. The objective weight of each photosynthetic parameter (W_j_) is derived by using the Criteria Importance Though Intercrieria Correlation (CRITIC) method, which reflects the relative importance by applying the comparative and conflict information among the indicators (Diakoulaki et al. 1995). Two calculation steps are used to evaluate the weights of each criterion using the CRITIC method.

Step 1 is to normalize the decision matrix using Equation (10).

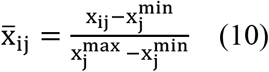

Step 2 is to calculate the weights for each criterion

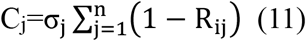

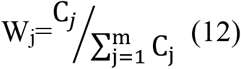

where, *σ*_*j*_ was the standard deviation of each criterion j, and R_ij_ is the linear correlation coefficient between the vectors X_i_ and X_j._ The comprehensive evaluation value of the effects of leaf anatomical traits on photosynthetic parameters were estimated from Equation (13).

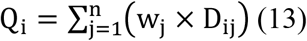

where D_ij_ was the correlation (direct effects, indirect effects, and Pearson’s correlation, respectively, respectively) between leaf anatomical trait i and photosynthetic parameter j . All statistical analyses were conducted with IBM SPSS Statistics 24.

## Results

Results showed the photosynthetic light-response curve (Fig.2) fits very well with the modified rectangular hyperbola model (R^2^=0.945 ± 0.006) described by Ye (2007) (Ye 2007). The *P*_n_ increased with *I* below the *I*_sat_. As *I* rise above the *I*_sat_, *P*_n_ decreases as *I* increase, implying a photoinhibition phenomenon. The calculated AQY would be 0.028 ± 0.003 if the Kok effect were ignored. This value was lower than *α* (0.031 ± 0.003) and the IQY (0.033 ± 0.003). The maximum *P*_n_ values calculated by the modified rectangular hyperbola model at *I*_sat_ of 621.762±37.358 μmol (photon) m^−2^s^−1^ was 5.131±0.478 μmol (CO_2_) m^−2^s^−1^ for coffee leaf. The values of *I*_c_ and *R*_D_ were 60.238 ± 11.332 μmol (photon) m^−2^s^−1^ and 0.930 ± 0.098 μmol (CO_2_) m^−2^s^−1^, respectively.

**Fig 2.**
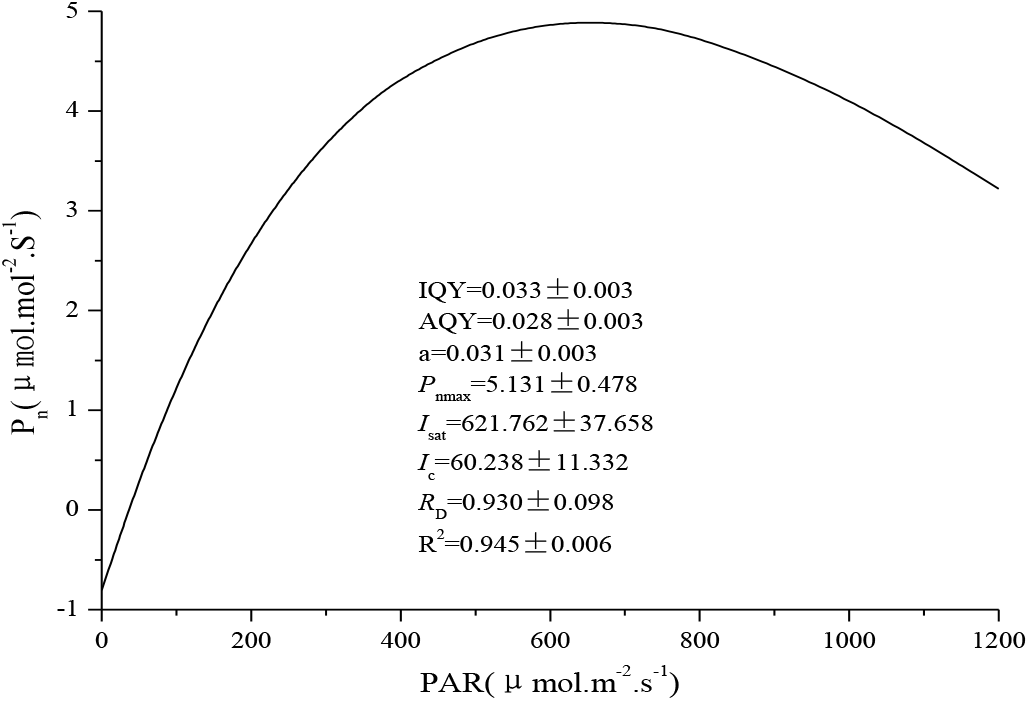
Photosynthetic light-response curve of coffee plants. Data shown in the figure indicate mean ± standard error. IQY – intrinsic quantum yield; AQY – apparent quantum yield; *α* – the absolute value of slope between *I* = 0 and *I* = *I*_c_; *P*_nmax_ – maximum photosynthetic rate [μmol (CO_2_) m^−2^ s^−1^]; *I*_sat_ – light saturation point [μmol (photon) m^−2^ s^−1^]; *I*_c_ –light compensation point [μmol (photon) m^−2^ s^−1^]; *R*_D_ – dark respiration [μmol (CO_2_) m^−2^ s^−1^].

The A/*C*_i_ curve was fitted with the interactive website (http://www.leafweb.ornl.gov) based on the FvCB model structure (Gu and Sun 2014). Figure 3 shows the response of A (net photosynthesis) to *C*_i_ for coffee leaves. The parameters of values for *V*_cmax_, *J*_max_, *T*_p_, *R*_d_, *Γ*\*, *g*_i_, and *K*_co_ were 61.749 ± 5.394, 63.165 ± 3.859, 5.316 ± 0.576, 3.225 ± 0.441, 3.309 ± 0.333, 1.101 ± 0.287, 103.096 ± 11.721, respectively. The transition between the Rubisco- and RuBP regeneration-limited states (*C*_i_CJ_) occurs around 20 Pa intercellular CO_2_ (Fig 3.).

**Fig 3.**
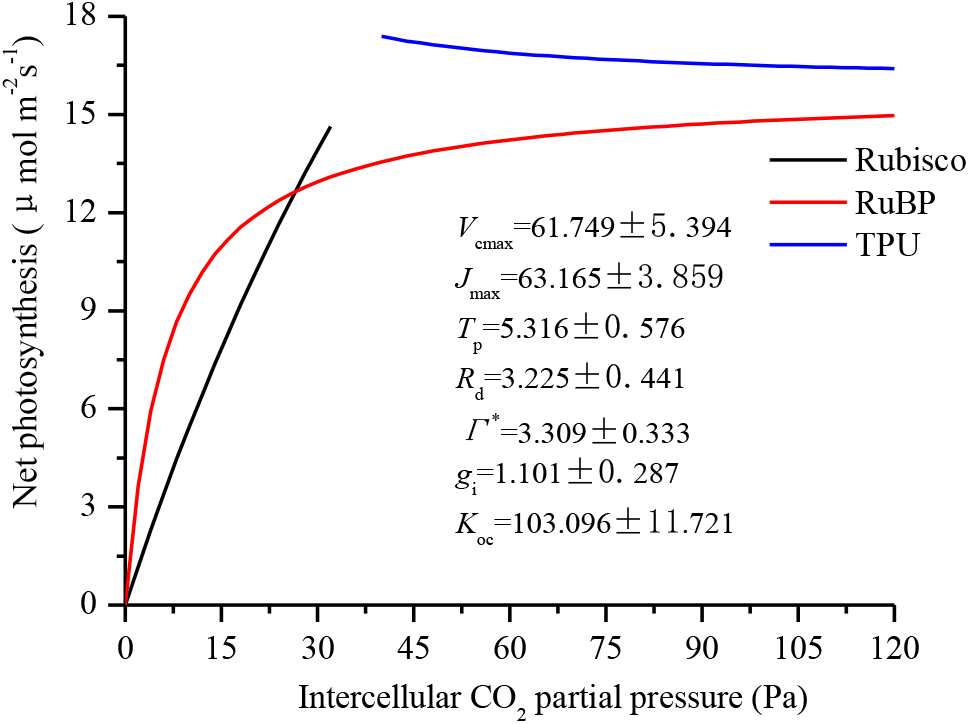
The A/C_i_ curves for the coffee plants. Data shown in the figure indicate mean ± stand error. *C*_i_–Intercellular CO_2_ concentration (μmol mol^−1^); *V*_cmax_–Maximal Rubisco carboxylation rate (μmol m^−2^ s^−1^); *J*_max_–Maximal electron transport rate (μmol m^−2^ s^−1^); *T*_p_–Rate of triose phosphate export from the chloroplast (μmol m^−2^ s^−1^);*R*_d_–Day respiration (μmol m^−2^ s^−1^); *Γ* *–CO_2_ compensation point in the absence of dark respiration (Pa); *g*_i_– Internal (mesophyll) conductance to CO_2_ transport (μmol m^−2^ s^−1^ Pa^-1^); *K*_*0*c_ –A composite parameter (Pa):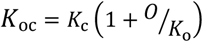.

Cross-sections observed with light microscope revealed that the coffee leaves were bifacial, with a typical well-differentiated single layer of elongated adaxial palisade and large abaxial spongy mesophyll cells. The large spongy mesophyll cells at the abaxial side comprised almost 60% of the blade thickness (Table 1). The variation of dimensions of the palisade tissue thickness (PT) is higher than in other tissues (CV=13.243%). The stomatal density (*d*) and total stomatal pore area (TSP) were 152.028 and 0.439, respectively (Table 1.).

**Table 1.** The anatomical of component tissues in Arabic coffee leaf.

| Leaf anatomy |  | Mean | SE | CV (%) | Proportion of Leaf Thickness (%) |
| --- | --- | --- | --- | --- | --- |
| Tissue | LT | 255.627 | 1.832 | 4.645 | 100.000 |
| Components<br>($\mu\text{m}$) | UCT | 5.497 | 0.042 | 4.962 | 2.154 |
|  | UET | 23.681 | 0.291 | 7.957 | 9.279 |
|  | PT | 51.014 | 1.042 | 13.243 | 19.913 |
|  | ST | 153.285 | 1.245 | 5.264 | 59.969 |
|  | LET | 17.908 | 0.195 | 7.056 | 7.023 |
|  | LCT | 4.243 | 0.040 | 6.121 | 1.662 |
| Stomatal<br>Characteristics | W ($\mu\text{m}$) | 8.252 | 0.048 | 3.733 | |
| | L ($\mu\text{m}$) | 27.874 | 0.208 | 4.839 | |
|  | *d* (No./mm<sup>2</sup>) | 152.028 | 1.942 | 8.277 |  |
|  | TSP | 0.439 | 0.006 | 8.886 |  |

The correlation between 19 physiological indexes and leaf anatomical traits was established by Pearson’s correlation analysis (Table 2). The UCT showed a negative correlation with the indexes of IQY, AQY, *α, I*_*sat*_, and *g*_i_ (*p*<0.05). The UET was positively related to AQY, *α*, and *P*_max_ (*p*<0.05) and negatively related to *I*_c_, *R*_d_, *C*_i_, and VPD (*p*<0.05) (Table 2). The relationship between PT and IQY, AQY, and α was significantly negative (*p*<0.05). There was a significant negative correlation between LET and *I*_c_ (*p*<0.05). The Pearson correlation coefficient between LCT and *I*_sat_ was significantly negative (*p*<0.05). The guard cell width (*w*) significantly affected *I*_c_ and *R*_d_ (*p*<0.05). The guard cell length (*l*) negatively correlated to IQY, AQY, *α*, and *I*_sat_, and positively correlated to *J*_max_ (*p*<0.05).A strongly positive (*p*<0.01) correlation relationship between stomatal density (*d*) and *J*_max_ was found in this study. However, Pearson Correlation study indicated that the effects of LT and ST do not significant affect the photosynthetic efficiency.

**Table 2.**
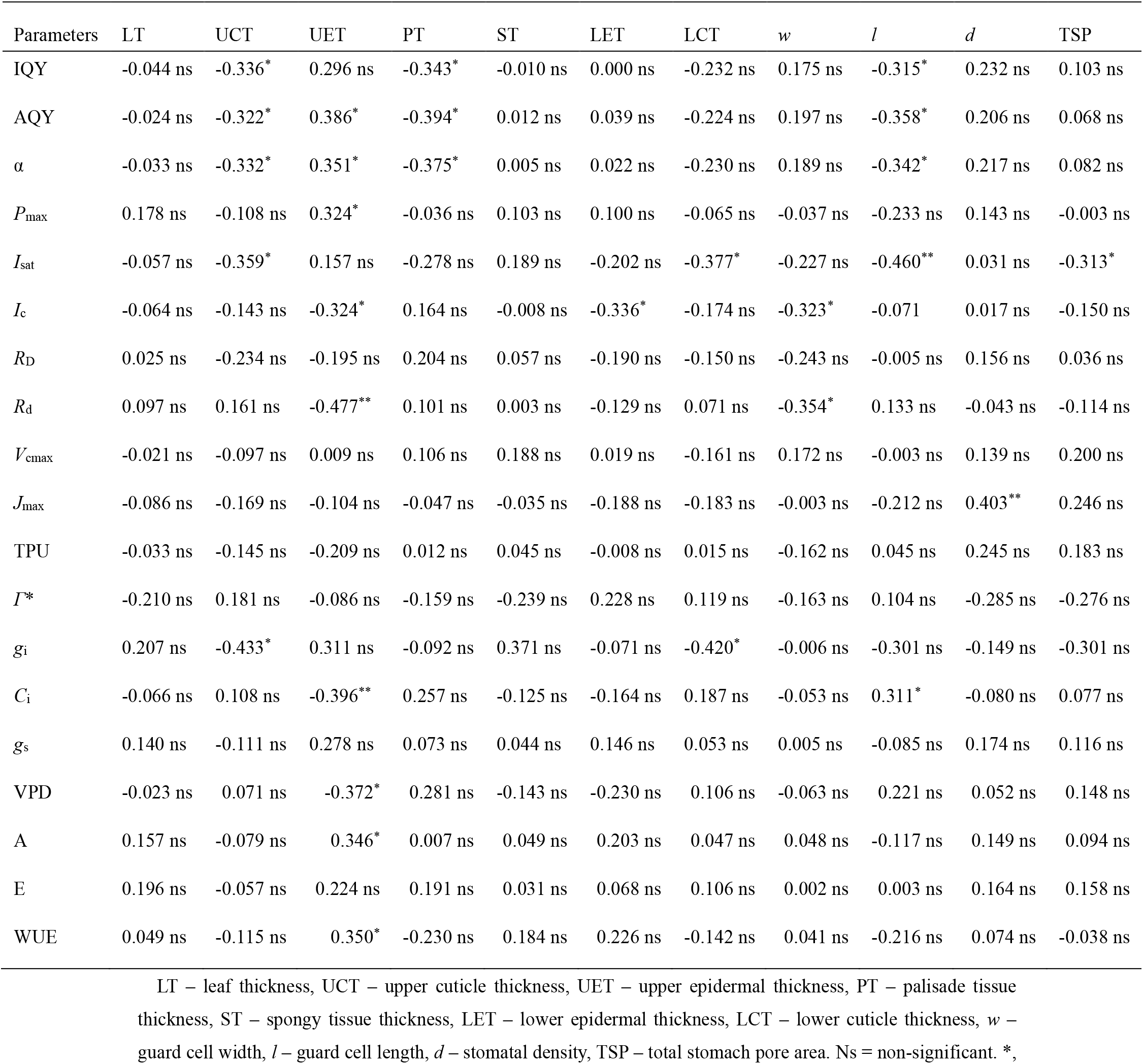
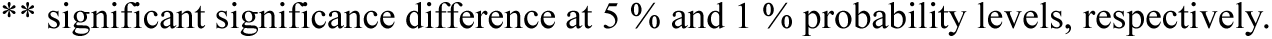
The correlation between leaf anatomical traits and photosynthetic parameters.

Path analysis was performed to reveal the direct and indirect effects of leaf anatomy traits on photosynthetic capacity. The results indicating the effect are shown in Figure 4. Although correlation analysis suggested that there were no significant correlations between LT and photosynthetic capacity variables (*p*>0.05), its direct effects on *g*_i_ (0.545), IQY (0.467), *R*_d_ (0.476), and *α* (0.420) were relatively high (Fig.4 (a)). In addition, there were high negative direct effects of LT on *I*_sat_ (-0.415). The path coefficient analysis revealed that UCT had a high magnitude negative direct effect on IQY (-0.686), *α* (-0.676), AQY (-0.652), *g*_s_ (-0.685), and TPU (-0.517). The direct effect of UET on *R*_d_ (-0.859) and TPU (-0.632) was highly negative. UET also had moderate positive direct effects on E (0.497), *P*_max_ (0.437), and A (0.423). PT exhibited an impressive negative direct effect on TPU (-0.474), AQY(-0.514), *α* (-0.506), and IQY (-0.487), while a high positive direct effect was shown between PT and E (0.445). ST presented a high positive direct effect on *I*_sat_ (0.630) and WUE (0.583). In addition, a similar high negative direct effect was also observed between ST and VPD (-0.579). LET presented high positive direct effects on WUE (0.482), *g*_i_ (0.416), TPU (0.462), and *V*_cmax_ (0.398). However, there was a relatively high negative direct effect between LET and *I*_c_ (-0.314), and VPD (-0.311). The LCT showed a negative direct effect with *V*_cmax_ (-0.539), *I*_c_ (-0.416), *R*_D_ (-0.411), *J*_max_ (-0.378), *I*_sat_ (-0.370), *g*_i_ (-0.340) and VPD (-0.328). The moderate positive direct effect of LCT on *Γ*\* (0.359) was observed. The stomatal density (*d*) exhibited large positive direct effects on *P*_max_ (0.893) *I*_sat_ (0.672), WUE (0.673), and *J*_max_ (0.482), and high negative direct effects on *C*_i_ (-0.772) and VPD (-0.576) were found. The negative direct effect of TSP on *I*_sat_ (-0.370) was very high. Besides, it also showed high direct effects on *g*_i_ (0.613), VPD (0.579), *P*_max_ (-0.519) and WUE (-0.496).

**Fig 4.**
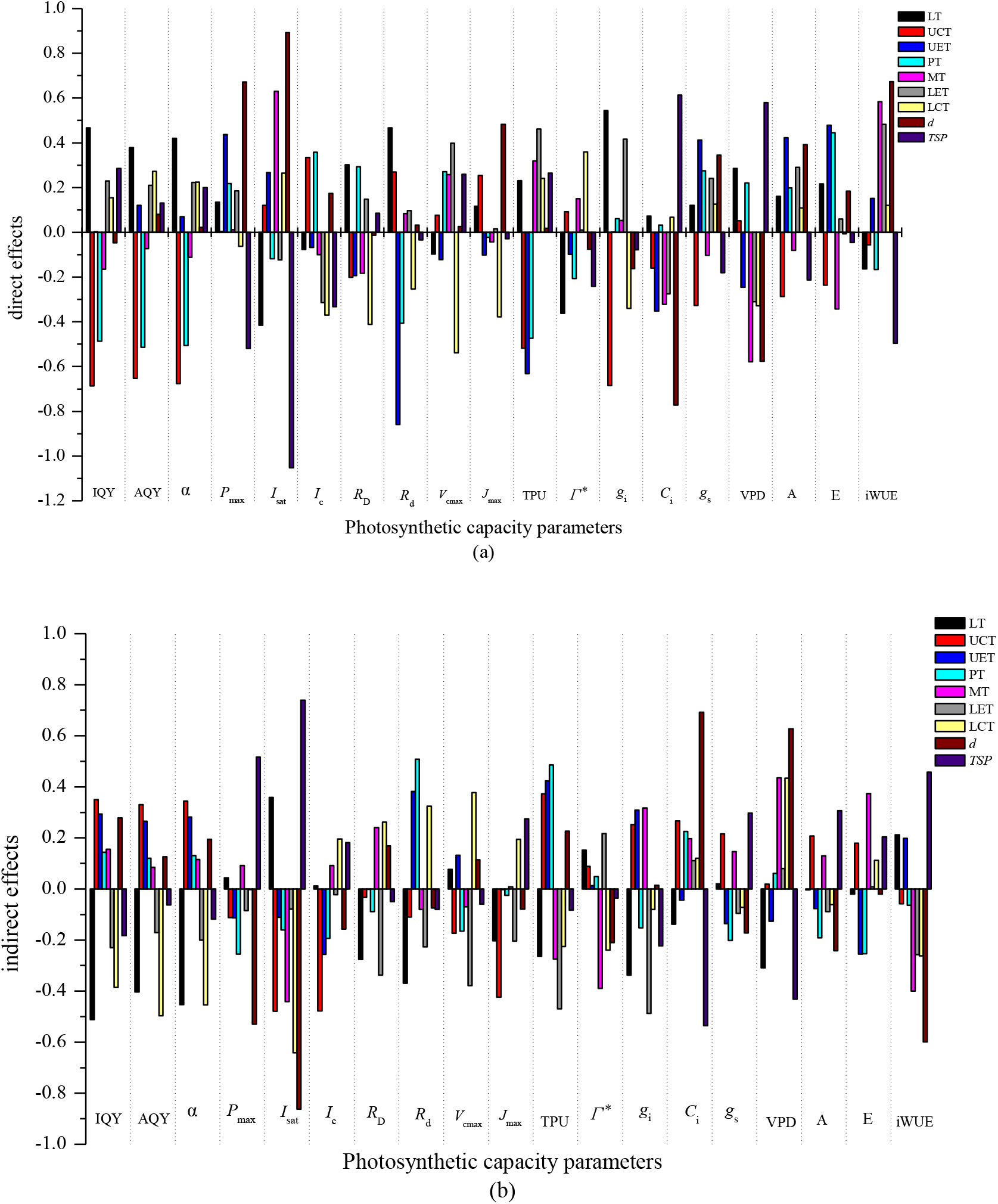
Direct (a) and indirect (b) effects of each leaf anatomical traits through other traits on photosynthetic parameters. LT – leaf thickness, UCT – upper cuticle thickness, UET – upper epidermal thickness, PT – palisade tissue thickness, ST – spongy tissue thickness, LET – lower epidermal thickness, LCT – lower cuticle thickness, *w* – guard cell width, *l* – guard cell length, *d* – stomatal density, TSP – total stomach pore area. The residual effect of IQY, AQY, *α, P*max, *I*_sat_, *I*_c_, *R*_D_, *R*_d_, *V*_cmax_, *J*_max_, TPU, *Γ*\*, *g*_i_, *C*_i_, *g*_s_, VPD, A, E, and WUE was 0.799,0.760,0.775, 0.849,0.737,0.859, 0.908,0.755,0.883,0.887, 0.860,0.877,0.644,0.817,0.858,0.821,0.840,0.858, and 0.801,respectively.

The estimates of each indirect effect of leaf anatomical trait on photosynthetic capacity via other leaf anatomical traits were presented in Figure 3 (b). According to this figure, there were highly negative indirect effects of LT on IQY (-0.5.11), *α* (-0.453), AQY (-0.403), *R*_d_ (-0.369), and *g*_i_ (-0.338). LT also had a high positive indirect effect on *I*_sat_ (0.358). UCT revealed a high indirect effect on *I*_sat_ (-0.479), *I*_c_ (-0.477), *J*_max_ (-0.423), TPU (0.372), IQY (0.350), AQY (0.330), *α* (0.344), and *g*_i_

(-0.319) through other anatomical traits. UET indirectly exerted relatively high positive effects on TPU (0.424) and *R*_d_ (0.382) via other anatomical traits. Similarly, PT exhibited relatively high positive indirect effects on *R*_d_ (0.508) and TPU (0.486) via other anatomical traits. The indirect effects of ST on *I*_sat_ (-0.441), *Γ*\* (-0.389), and WUE (-0.399) via other anatomical traits were negative and high. Also, relatively high positive indirect effects on *g*_i_ (0.318) and E (0.374) were detected for ST through other anatomical traits. LET revealed a positive indirect effect on *R*_D_ (-0.338), *V*_cmax_ (-0.379), TPU (-0.468), *g*_i_ (-0.487) through other anatomical traits. The indirect effects of LCT on IQY (-0.386), AQY (-0.497), *α* (-0.454), and *I*_sat_ (-0.642) were negative via other anatomical traits. However, there were indirect effects with a highly positive LCT on VPD (0.433) via other anatomical traits. Stomatal characteristics play a key role in determining photosynthesis. In this study, the *I*_sat_, WUE, and *P*_max_ were influenced indirectly with a relatively high negative magnitude by stomatal density (*d*) via other anatomical traits (-0.529, -0.862, and -0.599, respectively). There was also a relatively high positive indirect effect of stomatal density (*d*) on *C*_i_ (0.692), and VPD (0.628). The TSP presented a relatively high negative indirect effect on *C*_i_ (-0.536) and VPD (-0.432) through other anatomical traits. In addition, a highly positive indirect effect of TSP on *I*_sat_ (0.739), *P*_max_ (0.516), and WUE 0.458) was observed.

To identify the combined effect of leaf anatomical traits on photosynthetic parameters, a comprehensive evaluation was analyzed. Based on the evaluation values (Qi), the ranking of the comprehensive scores of relationships between photosynthetic capacity and leaf anatomical traits was shown in Table 2. The highest comprehensive score for direct effect was obtained by the LT, followed by LET, *d*, ST, PT, UET, TSP LCT, and UCT. For the comprehensive score of indirect effect of leaf anatomical traits on photosynthetic capacity, the UET held the highest comprehensive score, and UCT was the second, followed by ST, TSP, PT, *d*, LCT, LT, and LET. Similarly, results from the Pearson correlation analyzes of comprehensive assessment revealed that the first comprehensive assessment score was *d* and that the second assessment score was also ST. Then, the ranking order after the third was UET, LT, TSP, PT, LET, LCT, and UCT.

**Table 2.**
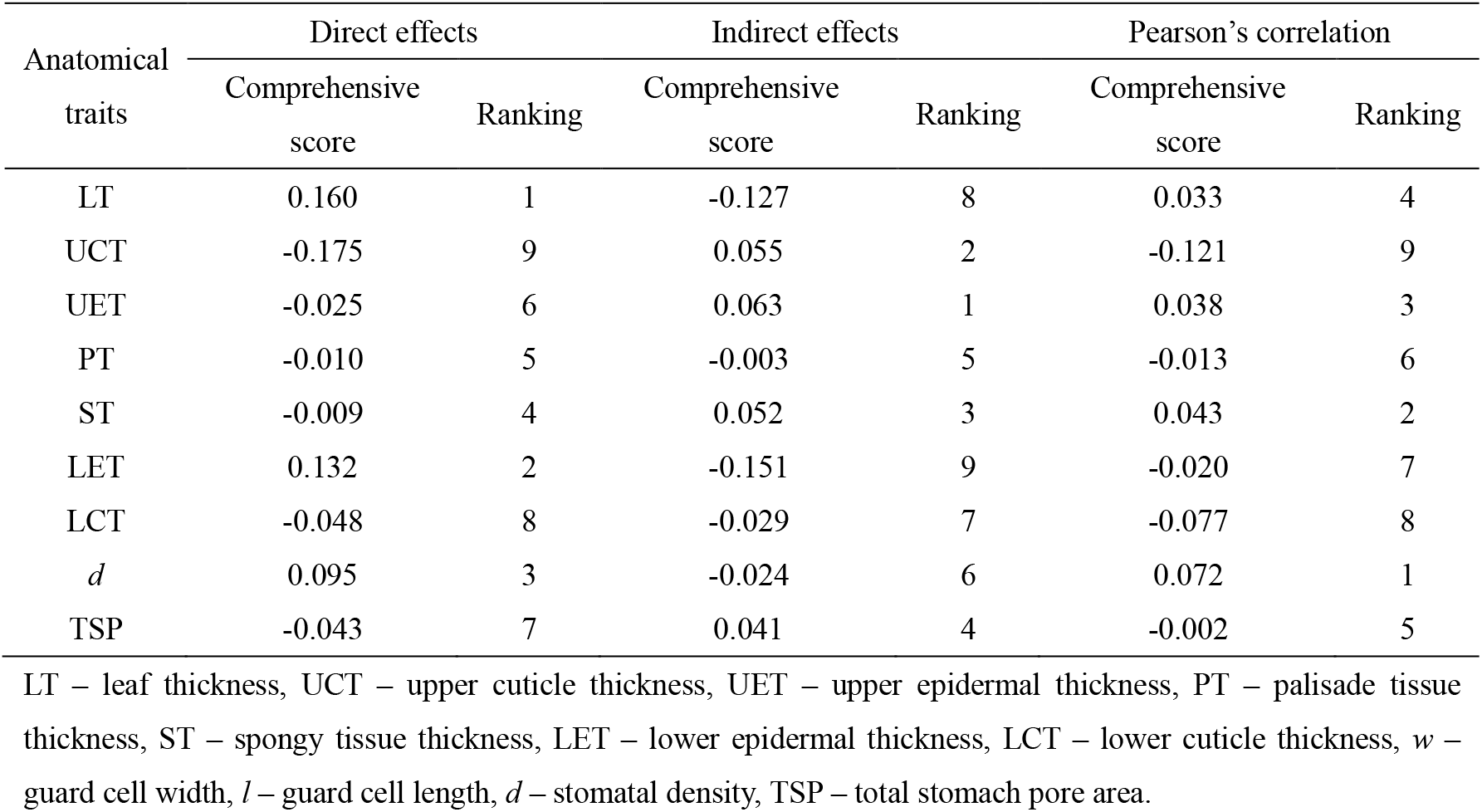
Comprehensive ordering of the effects of leaf anatomical traits on photosynthetic capacity with CRITIC method.

## Discussion

Several studies involving gas exchange measurements of coffee leaves have been performed. These studies have pointed out that the photosynthetic capacity of coffee leaves varied with a varying in environment (Ayalew 2018; DaMatta 2004; DaMatta and Ramalho 2006; Kumar and Tieszen 1980). For single coffee leaves, the saturating irradiance is between 300 and 700 μmol m^−2^.s^−1^, with shade leaves showing a lower value than lower sun leaves (DaMatta 2004; MATTA). This study found that the saturating irradiance of sun leaves was about 620 μmol photons m^-2^ s^-1^, which was consistent with the result of the above mention. Quantum yields revealed the relationship between a given light-dependent product and the number of absorbed photons (Taiz and Zeige 2006). Quantum yield is 0 when none of the light energy is used in photosynthesis. Quantum yield is 1 when all the absorbed light is used. Based on the results of YE (2007), the quantum yield decreased with *I* increasing (Ye 2007). Thus, the IQY was higher than AQY. In the past, there were several pertinent points about the maximal photosynthetic rates of coffee leaves. For example, Kumar and Tieszen (1980) and Cannell (1985) have pointed out that the maximal photosynthetic rates of sun leaves of coffee are around 7 μmol CO_2_ m^-2^ s^-1^ and 8.8 μmol CO_2_ m^-2^ s^-1^, respectively (Cannell 1985; Kumar and Tieszen 1980). However, Bote et al. (2018) reported that the maximal photosynthetic rates of sun leaves of coffee were lower than 7 μmol CO_2_ m^-2^ s^-1^ with a larger N supply (Bote et al. 2018). In this study, the maximal photosynthetic rates were lower than the observations of Kumar and Tieszen (1980) and Cannell (1985) and consistent with the results of Bote et al. (2018). For coffee leaves, light saturation would lead to an excess of electron transport capacity (Martins et al. 2014). Therefore, a higher *J*_max_ was observed under light saturation. In addition, the values of *V*_cmax_ and *J*_max_ were larger than the Martins et al. (2014) reported result.

The correlation analysis is a great utility method for understanding the relationships between variables. In this study, significant correlations between photosynthetic capacity and leaf anatomical traits were observed, indicating that leaf anatomical traits could affect photosynthetic capacity. Despite the correlation coefficients could provide a good way to understand the relationships between variables, the cause and effect relationships between variables couldn’t be estimated (Gonzalez-Paleo and Ravetta 2018). Path analysis, a method that investigates the causal relationship, gives detailed understanding of positive and negative correlations between traits (Gonzalez-Paleo and Ravetta 2018). The result of the path analysis was shown in figure 3, indicating that there were high-level direct and indirect effects of leaf anatomical traits on photosynthetic capacity. This may explain the importance of leaf structure in photosynthesis. However, the estimated residual effect in this study was high (0.661-0.908), indicating that additional characters which affect photosynthetic capacity are not included in the investigation. For leaf photosynthesis, the overall photosynthetic limitation can be partitioned into different components, such as stomatal, mesophyll, and biochemical limitations (Grassi and Magnani 2005). The stomatal and mesophyll limitations in coffee leaves accounted for 0.30 and 0.38, respectively (Martins et al. 2014). This is consistent with the results, where about 30% of the variability in photosynthetic capacity was contributed by the characters studied in the path analysis in this study. In order to quantitatively evaluate the effect of each structure of the leaf on photosynthesis, the comprehensive evaluation values are calculated by using the formula (13) introduced in the methods. The ordering Pearson’s correlation of comprehensive score showed that stomatal density (*d*) and spongy tissue thickness (ST) were the most important leaf anatomical traits for coffee leaf photosynthesis. For Arabica coffee, leaf photosynthesis is greatly limited by CO_2_ diffusion (DaMatta et al. 2007; Franck and Vaast 2009b). The CO_2_ diffusion from the air to the leaf mesophyll is mainly modulated by the stomatal conductance (g_s_), which is associated with stomatal density (*d*) (Lambers and Oliveira 2019; Rodrigues et al. 2018). According to the comprehensive score of indirect effect, the stomatal density (*d*) also had the highest comprehensive score. This indicates that stomatal density (*d*) is essential for coffee leaf photosynthesis. In fact, higher stomatal density will increase g_s_ by increasing the same total pore area (Rodrigues et al. 2018) and allow more CO_2_ diffusion. This may partly explain why the total stomach pore area (TSP) had higher comprehensive evaluation values in the effect of coffee leaf photosynthesis. Therefore, the result of a comprehensive evaluation revealed that Arabic coffee leaf of stomatal characteristics, such as guard cell width and length, stomatal density, and stomatal pore surface, plays a significant role in leaf photosynthetic capacity. This finding was also consistent with previous work, indicating stomatal characteristics of the leaf are closely associated with photosynthesis by controlling the water loss and CO_2_ uptake (Sakoda et al. 2020; Yin et al. 2020).

The leaf thickness was strongly positively correlated with the fraction of intercellular air space in the leaves (Niinemets 1999). The result obtained in this study showed that leaf thickness (LT) affects photosynthesis mainly through direct and indirect effects. Given that the CO_2_ diffusion in the gaseous phase is about three orders of magnitude larger than in the liquid (Oguchi et al. 2018). Thus, the diffusive resistance should not necessarily increase considerably with increasing leaf thickness (Smith et al. 1997). The result of direct effects of leaf thickness (LT) on photosynthetic capacity confirmed that increasing leaf thickness may increase *g*_i_. Moreover, due to the chloroplasts adhering to the inner surface of exposed mesophyll cell walls, thicker leaves provide space for more chloroplasts per unit leaf area (Lambers and Oliveira 2019) and possess more chloroplast surface area, which results in a substantial increase in carbon dioxide absorption because of a larger surface area for diffusion (Smith et al. 1997). The direct effects of the path analysis indicate that greater leaf thickness would facilitate CO_2_ diffusion in the mesophyll (*g*_i_). Furthermore, thicker leaves may have higher total contents of Rubisco and leaf N per unit area (Niinemets and Sack 2006), so it’s advantageous to realize high photosynthetic rates in highlight environments (Terashima et al. 2011). In this study, leaf thickness (LT) affects photosynthetic indexes that were IQY, *α*, AQY, *R*_d_, *g*_i_, and *I*_sat_, and, consequently, photosynthesis rate. Therefore, the result of a comprehensive evaluation revealed that leaf thickness (LT) was the most important role in the direct effect on leaf photosynthetic capacity.

The structure of the leaf cuticle consists of epicuticular wax, birefringent wax embedded in cutin, cutin matrix, and pectic substance (Lee and Graham 1986). The epicuticular wax can decrease light by scattering (Bote et al. 2018). Similar results have been found in this study where an increase in upper cuticle thickness (UCT) could result in a decrease in IQY, AQY, and *α*, which were related to light absorption in photosynthesis. The most cutin matrix in the cuticle is heterogeneous, and the areas of embedded waxes exhibit birefringence in polarized light (Lee and Graham 1986). This heterogeneous structure of the leaf cuticle may affect epidermal focusing, allowing actinic light to penetrate deeper into the mesophyll (Bote et al. 2018). These findings most likely explained the significant relationship between cuticle thickness and quantum yield of photosynthesis, such as IQY, AQY, and *α*. In addition, the cuticle thickness would increase resistance to CO_2_ diffusion (Miranda et al. 1981), and the negative relationships between UCT and LCT and photosynthetic parameters were also observed in this study.

The epidermal cell usually has a large transparent central vacuole and lacks mature chloroplasts, thus the light can directly pass from these cells into the mesophyll (Bote et al. 2018). However, these epidermal cells have various shapes, such as plano-convex, spherical, conical, or some other convex shape (Jackson et al. 2012). The curved outermost epidermal cell wall can affect the focal point within the leaf (Vogelmann et al. 1996). However, the result of Brodersen and Vogelmann (2007) showed that there was no relation between the convexity of epidermal cells and the absorptance of diffuse light (Brodersen and Vogelmann 2007). Although two contradictory hypotheses have been proposed concerning the impacts of epidermal cells on photosynthesis, the significant relationship between UET and photosynthetic capacity parameters in this study examined epidermal lens effects on many photosynthetic parameters and elucidated epidermal cells property plays an important role in photosynthetic capacity.

The columnar palisade tissue minimizes light scattering when the incident light collimated with the columnar palisade (Vogelmann and Martin 1993). As a result, palisade tissue allows large amounts of light to penetrate the chloroplasts within the leaf (Vogelmann and Martin 1993). The present study has revealed a remarkable relation between quantum yields and PT. However, the results of this study indicated that PT increase would directly decrease the IQY, AQY, and *J*_max_. This circumstance is very difficult to comprehend. Generally, palisade tissue has much more chloroplasts compared with other leaf tissue (Terashima and Inoue 1985). Thus, the photosynthetic activity of the palisade tissue would be higher. Indeed, the shape of cells in the leaves is strongly associated with photosynthetic performance by affecting the movement and distribution of chloroplasts (Gotoh et al. 2018). There was a great variety of dimensions of the palisade tissue thickness (PT) in this study. Coordinated regulation of leaf cell shape and chloroplast motion according to light conditions is essential for efficient leaf photosynthesis (Gotoh et al. 2018). This work could partially explain the result of a negative direct effect of PT on photosynthetic parameters.

The concentration of CO_2_ in the mesophyll is affected by air temperature, and each degree rise in temperature above 24°C results in a 20 ppm rise in CO _2_ concentration (Nunes et al. 1968). However, Khairi and Hall’s (1976) studies of citrus photosynthesis revealed that mesophyll conductance to CO2 decreased as temperature increased from 22 °C to around 40 °C (Khairi and Hall 1976). (Khairi and Hall 1976). In the analysis of the effects of leaf anatomical traits on photosynthetic capacity, the spongy tissue thickness (ST) had larger indirect effects and Pearson’s correlation comprehensive score, suggesting that spongy tissue thickness (ST) plays an important role in the photosynthetic process. Furthermore, Martins et al. (2014) demonstrated that mesophyll thickness is related to water flux and gas exchange per leaf area(Martins et al. 2014), which is consistent with the findings of this study. The spongy tissue thickness (ST) had direct effects on WUE and VPD. On the other hand, the irregularly shaped spongy cells alternating with air spaces increasese light absorption due to increase optical path length through a leaf (Oguchi et al. 2018), and this absorbed light will be advanced for photosynthesis (Brodersen and Vogelmann 2007; Cannell 1985). Therefore, the high direct spongy tissue thickness (ST) effects upon *I*_sat_ were also observed in this study. In fact, the increase in spongy tissue thickness (ST) would increase the surface area of chloroplasts exposed to intercellular airspace, which was associated with CO_2_ transfer conductance (Evans et al. 1994). Although there was little relationship between spongy tissue thickness (ST) and photosynthetic parameters, high direct spongy tissue thickness (ST) effects upon *I*_sat_, WUE, and VPD were observed.

Stomatal density (SD) is a factor associated with the photosynthetic rates of the leaf (Bondada et al. 1994). The CO_2_ diffusion conductance is influenced by the density and arrangement of the stomata (Oguchi et al. 2018). For the coffee leaf, the net carbon assimilation rate (A) is greatly governed by the diffusive conductance of CO_2_, thus the photosynthesis of the coffee leaf was largely limited by a diffusive factor (Araujo et al. 2008). Stomata have traditionally been thought to play an important role in controlling CO_2_ diffusion (Araujo et al. 2008) and were a key parameter in *C. arabica* photosynthesis (Ayalew 2018). Among the various stomatal characters, stomatal density plays a major role in gas exchange and photosynthesis (Pompelli et al. 2010). In this study, it is shown that stomatal density had the highest comprehensive scores compared with other anatomical traits on indirect effects and Pearson’s correlation comprehensive evaluation. In addition, structural modification of the stomata is one of the key factors affecting stomatal conductance (Taiz and Zeige 2006). The result of this study also showed that the guard cell length (*l*) significantly correlated positively with *C*_i_.

For light absorption, although the result of Martins et al. (2014) showed that the guard cell length did not change in response to light treatments (Martins et al. 2014), there was a significant correlation with IQY and AQY. The coordination between increased initiation of stomata cells and expansion of epidermal cells was observed in coffee leaves, implying that there was an optimization of the trade-off between transpiration costs and CO_2_ assimilation (Martins et al. 2014). In addition, the size and density of the stomata largely affect the diffusive conductance of the leaves to CO_2_, and the balance between the amount of CO_2_ required for photosynthesis and the level of water availability is also determined by the stomatal traits (Rodrigues et al. 2018). Thus, lower epidermal stomatal density had a significant correlation with photosynthetic parameters or direct effects and indirect effects on photosynthetic parameters. Moreover, these traits also had larger comprehensive scores in a comprehensive evaluation, which means that stomatal density was a key parameter in the regulation of photosynthesis of *C. arabica*.

## Conclusion

Many leaf anatomical traits significantly affect photosynthetic parameters. This result provided holistic relationships between leaf anatomical traits and photosynthetic parameters in coffee. It is tempting to suggest that these relationships might, to some extent, explain the photosynthetic behavior of coffee (*Coffea arabica* L.). The relationships between leaf anatomical traits and photosynthetic parameters emphasize the importance of leaf anatomy in determining photosynthesis at the tissue level. Moreover, these correlations between leaf anatomical traits and photosynthetic parameters were useful to explore the impact of different anatomical features on photosynthetic efficiency and accordingly provide some information to design leaf anatomy for enhanced photosynthetic efficiency. To a certain extent, path analysis and comprehensive evaluation could indicate an interrelationship between photosynthetic capacity and leaf anatomical traits. However, the causal interrelationship between photosynthetic parameters and leaf anatomical traits is less clear. Despite this, the result of the study also suggests that thinner leaves and higher stomatal densities might support a higher photosynthetic capacity. This result may be useful information for coffee breeding programs and plantation management.

## Author contributions

Z.X. and G.M. conceptualization and research design; X.H. and J.L. project administration; M.Z.and L.S. investigation and fieldwork, Z.X. analyzed the data and wrote the manuscript; H.M. and H.Z. resources; all the authors read and edited the manuscript, and approved the final version.

## Funding

This research was funded by the Basic Research Project of the Dehong Tropical Agriculture Research Institute of Yunnan (DTARI-JJ2020-02) and the DR155 Coffee Variety Industrialization Technology Integration and Application Demonstration (202003AD150006), and the National Tropical Plants Germplasm Resource Center.

## Conflict of interest statement

All authors certify that they have no affiliations with or involvement in any organization or entity with any financial interest or non-financial interest in the subject matter or materials discussed in this manuscript.

## Notes

### Competing Interest Statement

The authors have declared no competing interest.

### Summary of Updates

Addition: Fig.1 Path diagram for the relationship between leaf anatomical traits and photosynthetic parameters. Version of Fig. 3: Direct (a) and indirect (b) effects of each leaf's anatomical traits through other traits on photosynthetic parameters. Version of Table 2. : Comprehensive ordering of the effects of leaf anatomical traits on photosynthetic capacity with the CRITIC method.

